# CRISPR/Cas9-mediated gene editing induces neurological recovery in an A53T-SNCA overexpression rat model of Parkinson’s disease

**DOI:** 10.1101/2020.08.27.269522

**Authors:** Hyung Ho Yoon, Sunghyeok Ye, Sunhwa Lim, Seung Eun Lee, Soo-Jin Oh, Ara Jo, Hawon Lee, Na-Rae Kim, Kyoungmi Kim, Bum-Joon Kim, C. Justin Lee, Min-Ho Nam, Junseok W. Hur, Sang Ryong Jeon

## Abstract

To date, no publicly available disease-modifying therapy for Parkinson’s disease has been developed. This can be partly attributed to the absence of techniques for *in vivo* deletion of the SNCA gene (encoding α-synuclein), which is one of the key players in Parkinson’s disease pathology. In particular, A53T-mutated SNCA (A53T-SNCA) is one of the most studied familial pathologic mutations in Parkinson’s disease. Here we utilized a recently discovered genome editing technique, CRISPR/Cas9, to delete A53T-SNCA *in vitro* and *in vivo*. Among various CRISPR/Cas9 systems, SaCas9-KKH with a single guide RNA (sgRNA) targeting A53T-SNCA was packaged into adeno-associated virus. Adeno-associated virus carrying SaCas9-KKH significantly reduced A53T-SNCA levels in A53T-SNCA-overexpressed HEK293T cells, without off-target effects on wild-type SNCA. Furthermore, we tested the technique’s *in vivo* therapeutic potential in a viral A53T-SNCA overexpression rat model of Parkinson’s disease. Gene deletion of A53T-SNCA significantly prevented the overexpression of α-synuclein, dopaminergic neurodegeneration, and parkinsonian motor symptoms, whereas a negative control without sgRNA did not. Our findings propose CRISPR/Cas9 system as a potential therapeutic tool for A53T-SNCA familial Parkinson’s disease.

## Introduction

Parkinson’s disease (PD) is the second-most common neurodegenerative disease characterized by dopaminergic depletion in the nigrostriatal pathway and the presence of α-synuclein-containing Lewy bodies.^1^ The characteristic motor symptoms of PD are tremor, rigidity, and bradykinesia. The most common medication used for treating PD is L-3,4-dihydroxyphenylalanine (L-DOPA, also known as levodopa), which is a blood-brain barrier-permeable dopamine precursor.^2^ Even though levodopa is effective in alleviating the motor symptoms of PD, long-term administration of this drug causes several severe adverse effects including L-DOPA-induced dyskinesia.^3, 4^ More importantly, no available medication including L-DOPA can stop or modify the progression of PD.^4^

Several genetic studies have been aimed at identifying PD-related genes to determine the precise mechanism of PD. In particular, genome-wide association studies (GWAS) have identified variants of many candidate PD genes, such as variations of the SNCA gene encoding the α-synuclein protein.^5–7^ SNCA has been not only well documented to be engaged in rare familial forms of PD with its mutations and copy number variants,^8, 9^ but is also considered to be a major risk factor for sporadic PD because of its polymorphisms.^6^ Among the various mutations of SNCA, the missense mutation of Ala53Thr (A53T) in SNCA is one of the most well-known risk factors for early-onset PD,^8^ which acts by accelerating α-synuclein aggregation.^10^ Therefore, it is plausible that gene editing of the A53T-mutated SNCA (A53T-SNCA) gene could be a potent therapeutic approach for modifying disease progression in PD.

To develop a therapeutic approach for PD, two kinds of animal models are widely used: chemically-induced and genetic models.^11–13^ The most widely used chemically-induced models of PD are 1-methyl-4-phenyl-1,2,3,6-tetrahydropyridine (MPTP) and 6-hydroxydopamine (6-OHDA) models. Although these toxin-induced models show nigrostriatal tyrosine hydroxylase (TH) loss, decrease in striatal dopamine (DA) levels, and motor deficits,^14^ they do not resemble the main pathology of human PD, which is α-synuclein aggregation and disease progression.^15^ On the other hand, genetic models of PD have focused on mutating or knocking out genes known to cause familial PD. In particular, the genetic mutation of A53T-SNCA is one of the most widely used models. In the present study, we virally overexpressed A53T-SNCA in the substantia nigra pars compacta (SNpc), which has been well documented to exhibit not only nigrostriatal TH loss but also α-synuclein aggregation and progressive dopaminergic (DAergic) neuronal degeneration within a few weeks, leading to parkinsonian motor deficits.^16–18^

In 2013, Clustered Regularly Interspaced Short Palindromic Repeats (CRISPR) was first introduced as a tool for gene therapy in mammalian cells.^19, 20^ Despite accumulated evidence of its potential therapeutic effects *in vitro*, the lack of an adequate delivery method has hindered the development of gene therapy using CRISPR.^21, 22^ The adeno-associated virus (AAV), a clinically promising delivery vehicle of engineered genes, has been considered an attractive vehicle because of its low immunogenic response, low host genome integration, low oncogenic risk, and a high range of serotype specificity.^23–25^ However, its restrictive cargo size (~4.5 kb) has diminished the attraction of AAVs as a CRISPR vehicle.^26^ The most commonly used CRISPR-associated protein 9 (Cas9), *Streptococcus pyogenes* Cas9 (SpCas9), which shows high efficiency in gene editing, cannot be packed into AAV with other essential elements, because of its large size (4.2 kb). Recently, *Staphylococcus aureus* Cas9 (SaCas9), which is smaller (3.2 kb) than SpCas9, was developed and successfully packed in a single AAV with sgRNA and other expression elements.^27^ Subsequently, *in vivo* gene therapy using AAV-SaCas9 was reported to be successful in a mouse model of Duchenne muscular dystrophy.^28–30^ However, the complex sequence of the proto-spacer adjacent motif (PAM) of SaCas9 (5’-NNGRRT-3’) has restricted its use in various targets. Even though this limitation of SaCas9 was recently resolved in part by the development of SaCas9-KKH by random mutagenesis in the PAM recognition position (5’-NNNRRT-3’),^31^ the SaCas9 family has exhibited low efficiency in gene editing, compared to SpCas9. For these reasons, although the use of CRISPR has been underway to target a variety of genetic diseases in animal models, only a few successful examples have so far been reported.^32^

In this study, we constructed a SaCas9-KKH vector with sgRNA targeting A53T-SNCA and directly delivered it into the SNpc through AAVs. We further tested the therapeutic effect of this strategy in PD motor symptoms in an animal model of PD with virally overexpressed A53T-SNCA (A53T rat model).^18^ The CRISPR-mediated A53T-SNCA gene deletion caused a significant prevention of dopaminergic neuronal loss and a dramatic behavioral recovery. These findings suggest that CRISPR-mediated genome editing therapy could be a potent therapeutic strategy against A53T-SNCA familial PD.

## Results

### Construction of SaCas9-KKH with sgRNA specifically targeting A53T-SNCA

Using the ability to induce a double strand breakage at a target region, we hypothesized that CRISPR/Cas9 could be used to disrupt expression of the A53T-SNCA gene, which is known to be a major genetic cause of PD.^8^ To address this hypothesis, we sought to design sgRNA that specifically targeted the 53rd amino acid threonine of the A53T-SNCA gene, thereby avoiding wild-type SNCA (WT-SNCA) gene editing. For *in vivo* delivery of CRISPR/Cas9 through AAVs, we determined to use SaCas9 whose size is ~1 kb smaller than that of SpCas9 because of the limited cargo size of AAVs. We tried to design a proper sequence of sgRNA that simultaneously targets the 53rd amino acid threonine of A53T-SNCA and matches with the SaCas9 PAM sequence (5’-NNGRRT-3’). However, we could not find any matching sequence of sgRNA that met both criteria. Therefore, we considered the possible use of SaCas9-KKH whose PAM sequence is simpler than that of the original SaCas9. Finally, we designed a proper sgRNA sequence that specifically targeted A53T-SNCA (mismatch for WT-SNCA) and matched with the PAM sequence of SaCas9-KKH (5’-NNNRRT-3’) (Figure 1A). For the *in vitro* cleavage assay, we purified the SaCas9-KKH protein and synthesized the designed sgRNA. Subsequently, the SaCas9-KKH and A53T-SNCA-targeting sgRNA were packaged together into a single AAV reporter plasmid with a CMV promotor (pAAV-SaCas9-KKH-sgRNA) for the *in vivo* assay (Figure 1B). For negative control experiments, SaCas9-KKH without sgRNA was also cloned into the same single AAV reporter plasmid (pAAV-SaCas9-KKH) (Figure 1B).

**Figure 1.**
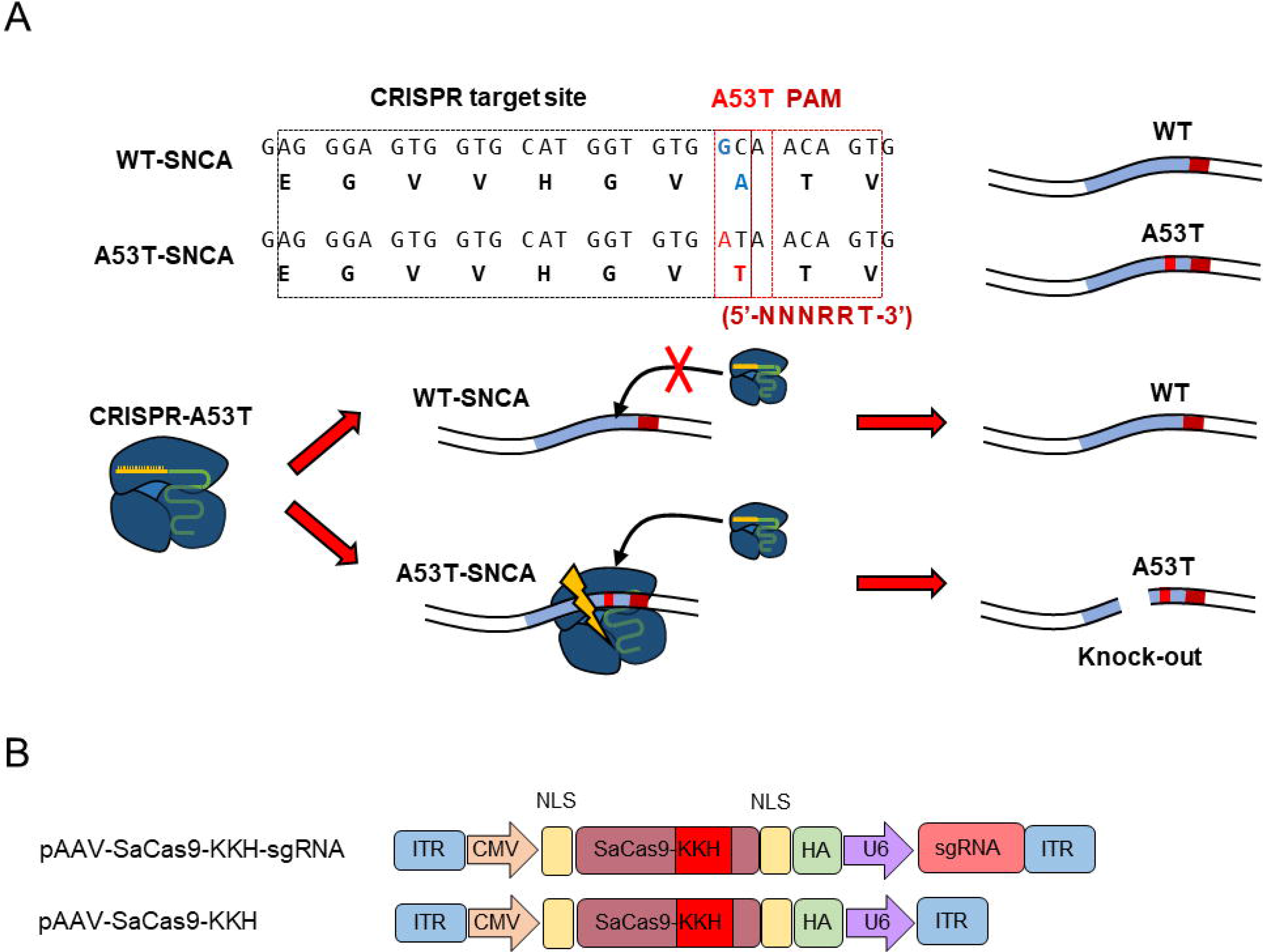
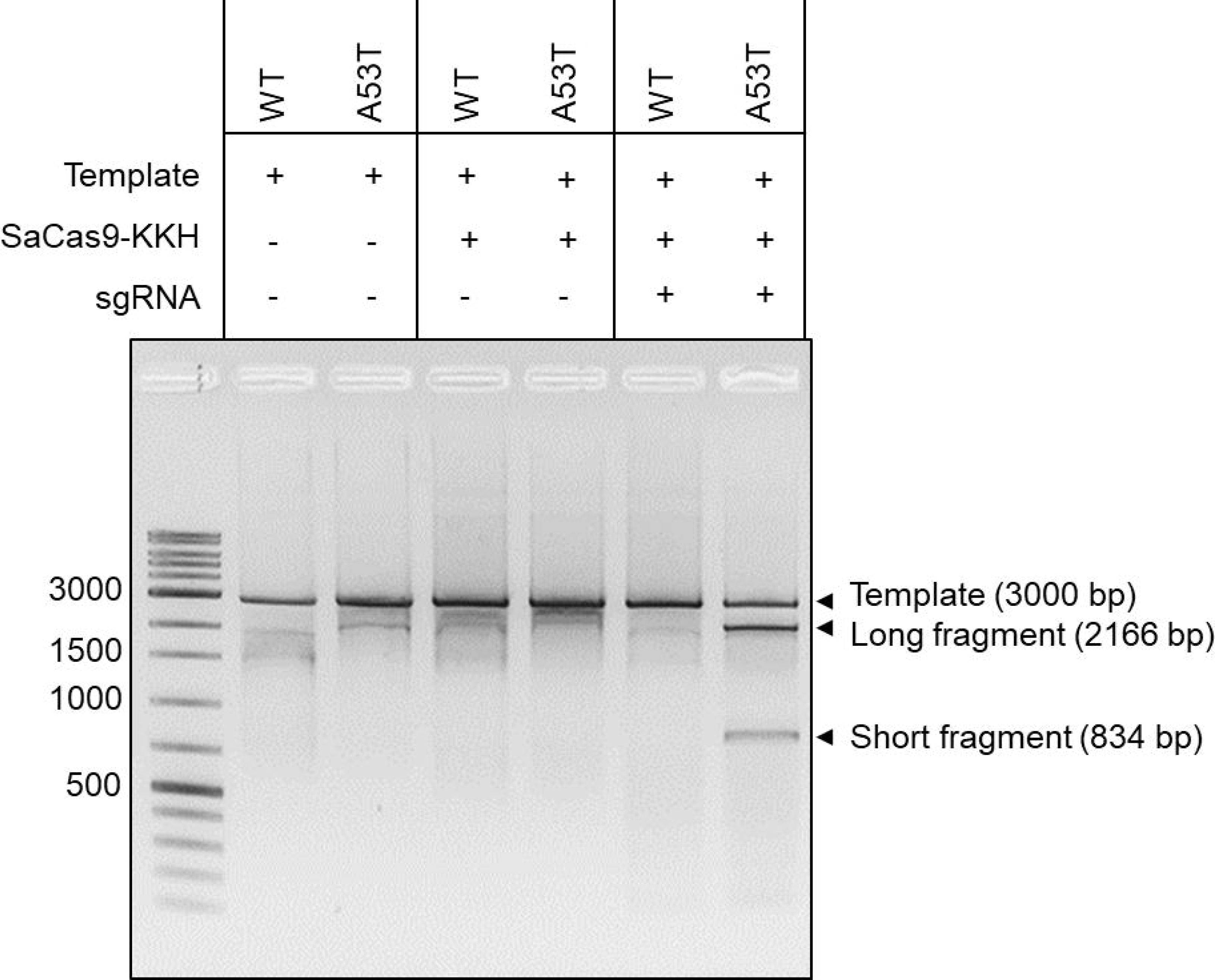
Schematic diagram of CRISPR and vector design. (A) Sequence information of sgRNA targeting the SNCA A53T region. To make a double strand breakage at the A53T sequence, we designed and cloned SaCas9-KKH with sgRNA in a single P3 vector that met the condition of PAM and proper breakage location. This system makes a double strand breakage at A53T-SNCA only without WT-SNCA disruption. (B) SaCas9-KKH and the A53T-SNCA-targeting sgRNA were packaged together into a single AAV reporter plasmid with the CMV promotor (pAAV-SaCas9-KKH-sgRNA). For negative control experiments, SaCas9-KKH without sgRNA was also cloned into the same single AAV reporter plasmid (pAAV-SaCas9-KKH).

### *In vitro* cleavage assay reveals CRISPR successfully cleaves A53T-SNCA without WT-SNCA disruption

The sgRNA and SaCas9-KKH protein (ribonucleoprotein; RNP) were incubated with WT-SNCA or A53T-SNCA templates. The A53T-SNCA template was cleaved and it showed two fragmented bands; however, the WT-SNCA template showed no fragmentation (Figure 2). These results indicate that our CRISPR system worked correctly at A53T-SNCA without any off-target event at WT-SNCA, in *in vitro* conditions.

**Figure 2.**
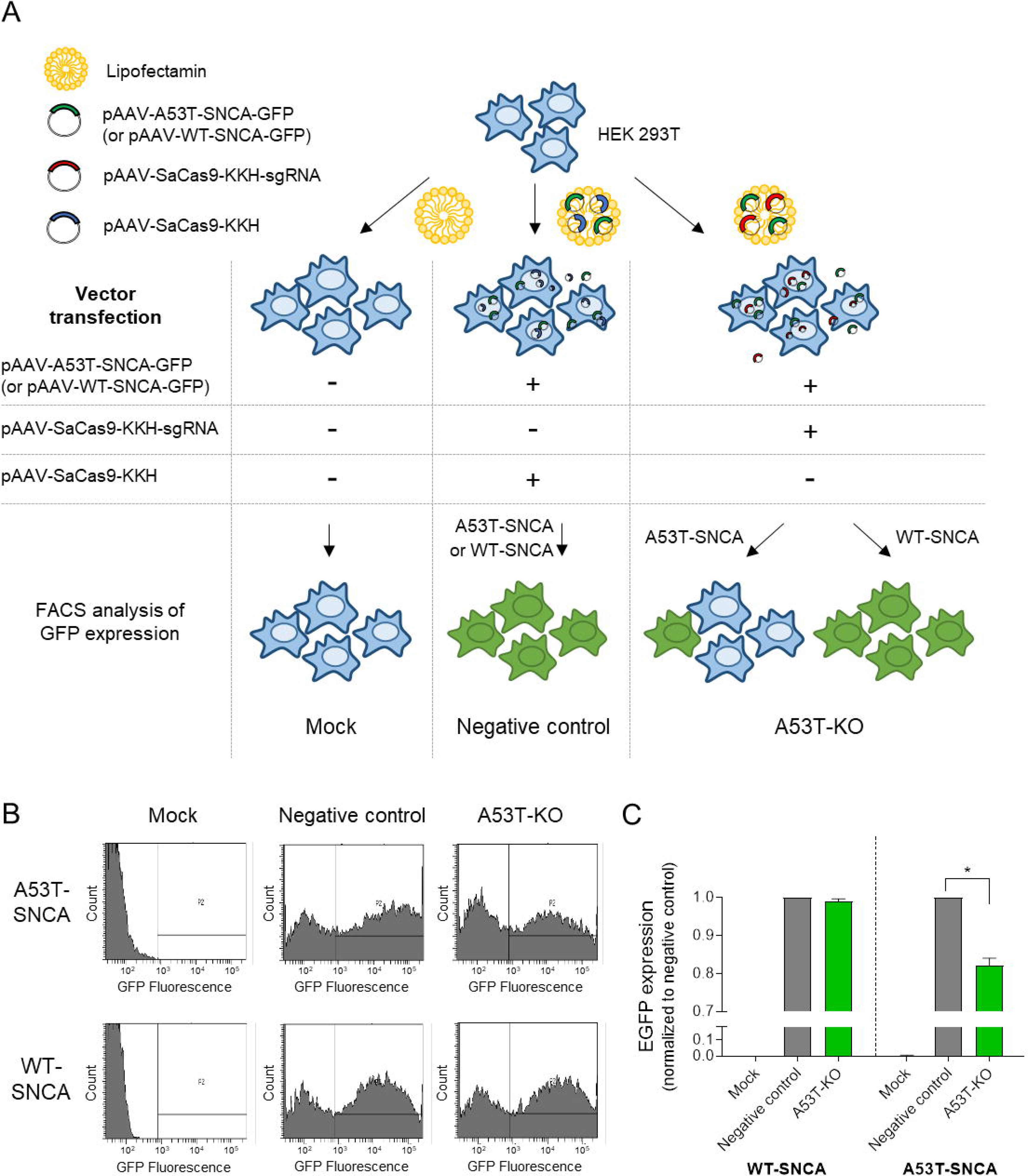
*In vitro* cleavage assay with WT-SNCA and A53T-SNCA. Templates containing WT-SNCA and A53T-SNCA sequences were incubated with SaCas9-KKH and sgRNA, and then analyzed by agarose gel electrophoresis. In the full CRISPR system, the A53T-SNCA template broke into two fragments, whereas the WT-SNCA template showed no change.

### CRISPR successfully disrupts A53T-SNCA without WT-SNCA off-targeting in cells

To validate the efficiency of pAAV-SaCas9-KKH-sgRNA in gene editing of A53T-SNCA in-cell, we prepared the HEK293T cell line transfected with A53T-SNCA with green fluorescent protein (GFP) using an AAV vector (pAAV-A53T-SNCA-GFP). To ensure the absence of any undesirable off-target issues on WT-SNCA, we also constructed a WT-SNCA-GFP vector and transfected it into HEK293T cells (pAAV-WT-SNCA-GFP). Simultaneously, we transfected pAAV-SaCas9-KKH-sgRNA and pAAV-SaCas9-KKH into HEK293T cells overexpressing A53T-SNCA-GFP and WT-SNCA-GFP, respectively. In parallel, using a knock-out (KO) validation assay, we confirmed that both CRISPRs (with and without sgRNA) were properly expressed in HEK293T cells after transfection, by quantitative real-time polymerase chain reactions (qPCR) and western blotting (Figure S1). Finally, we validated whether the CRISPR sufficiently disrupted A53T-SNCA expression, without WT-SNCA off-target events, by assessing the expression of GFP with fluorescence-activated cell sorting (FACS) analysis. Because SNCA was directly conjugated with GFP, the level of GFP expression indicated the level of SNCA expression. We found no GFP-expressing cells in mock groups; however, distinct populations of GFP-expressing cells were detected in both negative control groups (A53T-SNCA and WT-SNCA), which harbor CRISPR without sgRNA. In A53T KO groups (transfected with both A53T (or WT)-SNCA-GFP and CRISPR with A53T-targeting sgRNA), the pAAV-A53T-SNCA-GFP group showed a significantly reduced population of GFP-expressing cells. However, in the pAAV-WT-GFP group, the GFP-expressing cell population showed no change. These findings indicate that pAAV-SaCas9-KKH-sgRNA significantly and selectively disrupts gene expression of A53T-SNCA, and not of WT-SNCA, implying that the CRISPR targeting A53T-SNCA does not induce off-target effects on WT-SNCA (Figure 3).

**Figure 3.**
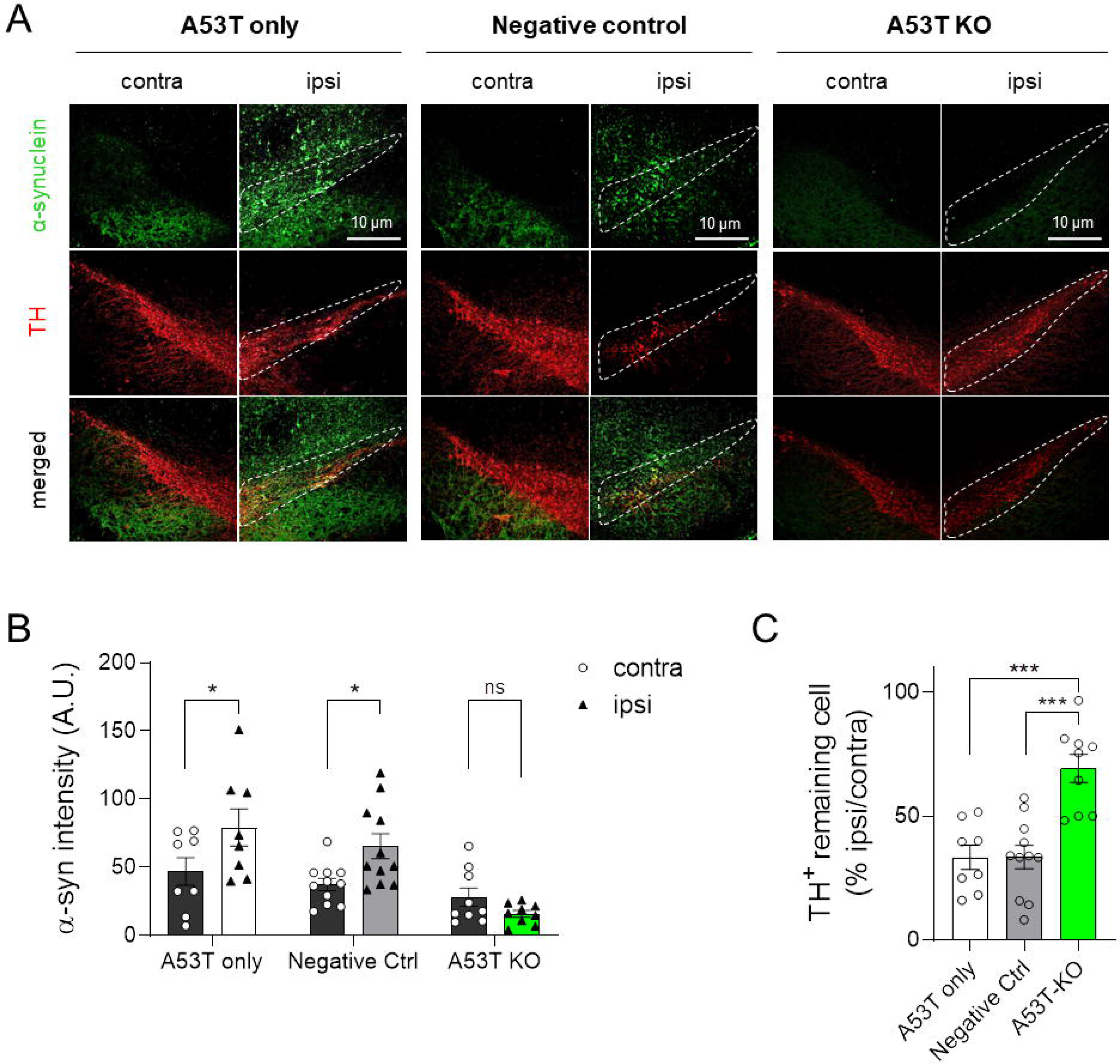
CRISPR/SaCas9-KKH significantly and selectively disrupts the expression of A53T-SNCA. (A) The HEK293T cell line was used for the CRISPR cleavage assay in-cell. Identical experiments were held parallel with A53T-SNCA and WT-SNCA, respectively. We divided the cells into three groups; i) no vectors, ii) A53T-SNCA (or WT-SNCA)+SaCas9-KKH, and iii) A53T-SNCA (or WT-SNCA)+SaCas9-KKH-sgRNA, with electroporation performed under the same condition with a Neon electroporator. After a 3-day incubation period, the GFP expression pattern was analyzed with FACS. (B) The first group (no vectors) shows no GFP expression. Negative control group shows high GFP expression level. For the A53T KO groups, A53T-SNCA-GFP with SaCas9-KKH-sgRNA group shows a clear frame shift to the left, which means that GFP expression has reduced significantly. However, the experiment with WT-SNCA shows there was no change in GFP expression. These results indicate that SNCA-GFP vectors produce target proteins properly and that the SaCas9-KKH-sgRNA vector also works well to make a double strand breakage at the intracellular target site (A53T-SNCA) without any off-target effect on WT-SNCA.

### AAV-CRISPR-A53T prevents α-synuclein overexpression and TH loss

Next, we evaluated whether the SaCas9-KKH nuclease can reduce the A53T-SNCA protein *in vivo* in the A53T rat model. We injected AAV-A53T-SNCA simultaneously with AAV-SaCas9-KKH-sgRNA or AAV-SaCas9-KKH into unilateral SNpc. Viral injection of AAV-A53T-SNCA is reported to induce significant parkinsonian motor deficits in rodents with a significant decrease in TH levels, the key dopamine-synthesizing enzyme, within 3 to 4 weeks.^18^ Therefore, to evaluate the expression level of the SNCA protein, we performed immunohistochemistry with an antibody against α-synuclein in the fourth week after virus injection. As a prerequisite, we confirmed by immunohistochemistry using an HA tag that the CRISPR protein was properly expressed in ipsilateral SNpc. (Figure S2). Subsequently, we found that α-synuclein levels were significantly increased in the ipsilateral SNpc compared to contralateral SNpc of the A53T rat model (Figure 4A). Intriguingly, we also found that AAV-SaCas9-KKH-sgRNA injection significantly reduced α-synuclein levels in the ipsilateral SNpc, whereas AAV-SaCas9-KKH without sgRNA did not (Figure 4B, C). These findings indicate that gene editing of A53T-SNCA through the CRISPR/Cas9 technique significantly blocks overexpression of α-synuclein *in vivo*.

**Figure 4.**
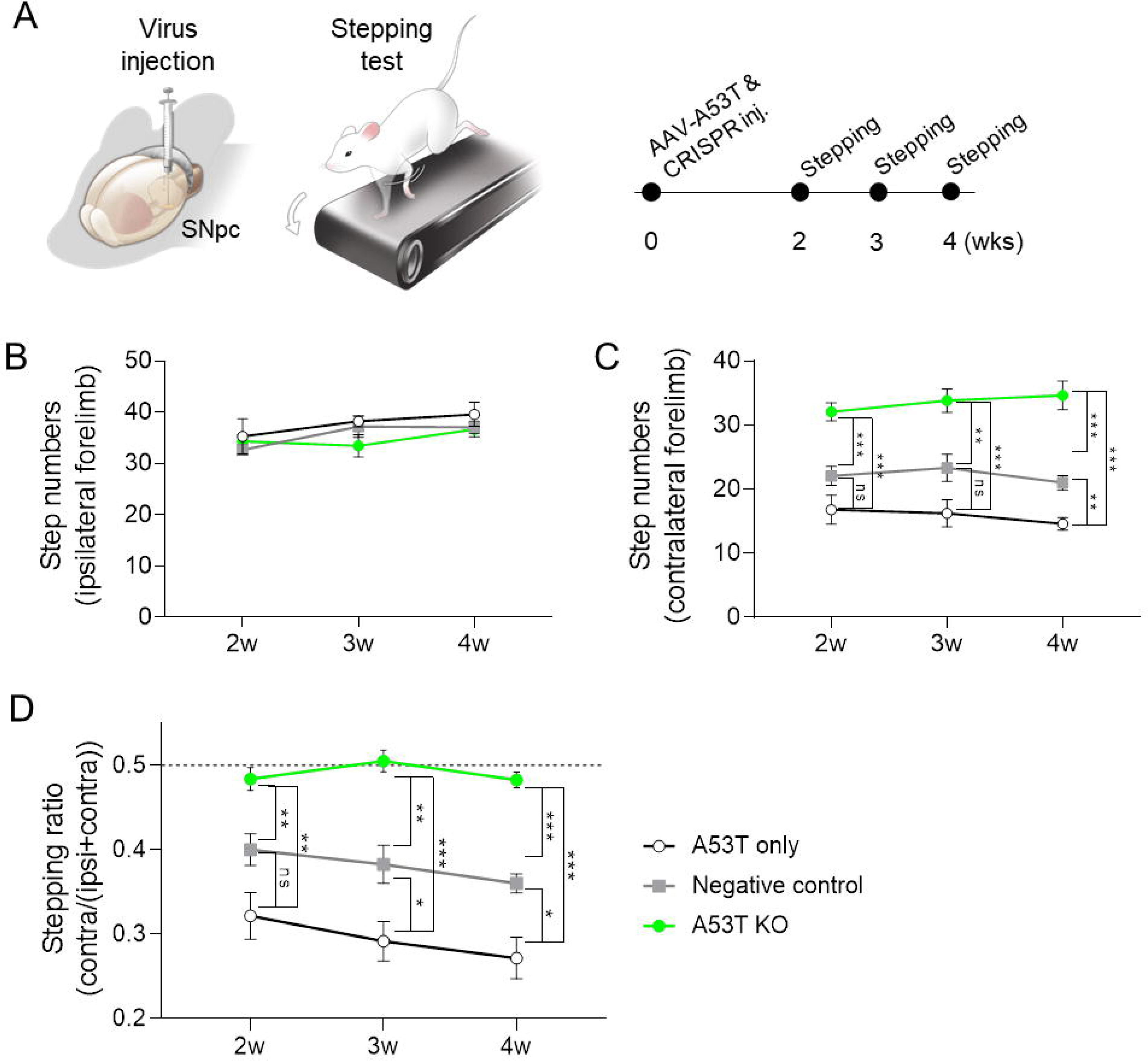
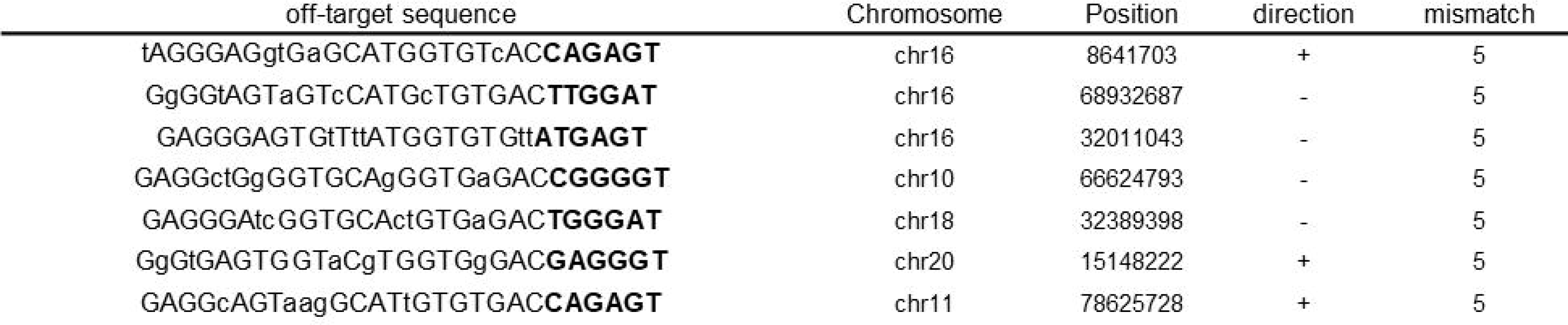
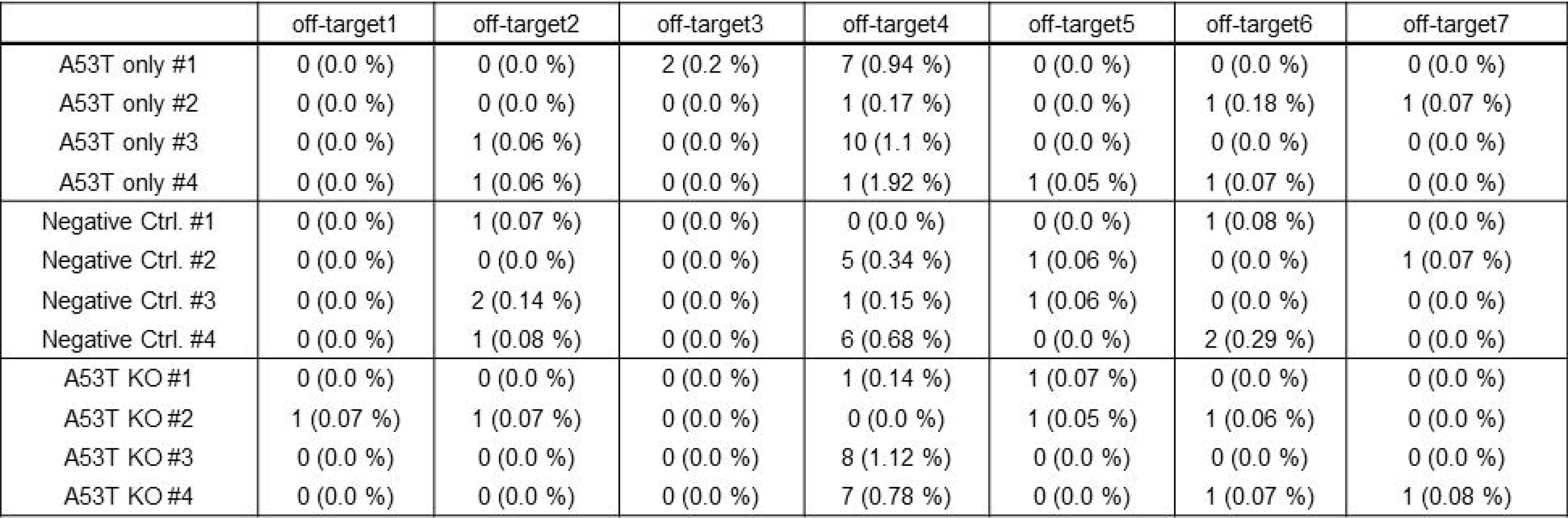
CRISPR-mediated A53T deletion significantly prevents A53T-SNCA-induced TH loss. (A) Representative immunofluorescence images of α-synuclein (green) and TH (red) expression in substantia nigra sections from A53T-SNCA-induced PD rat injected with PBS (A53T only), AAV-SaCas9-KKH without sgRNA (negative control), or AAV-SaCas9-KKH-sgRNA (A53T KO). (B) Quantification of α-synuclein immunoreactivity in SNpc (n = 4 rats per group). (C) Quantification of the remaining portion of TH-positive dopaminergic neurons in ipsilateral SNpc, compared to contralateral SNpc. Mean ± SEM. *P < 0.05; **P < 0.01; ***P < 0.001; ns, non-significant. Statistical significance was determined by the two-way ANOVA with Bonferroni’s multiple comparison test (B) or one-way ANOVA with Tukey’s multiple comparison test (C).

We also tested if the CRISPR/Cas9-mediated blockade of α-synuclein expression is associated with dopaminergic neurodegeneration in the A53T rat model. We performed immunohistochemistry with an antibody against TH, the marker for dopaminergic neurons. We found that only 32.5% of TH-positive neurons remained in the ipsilateral SNpc of the PD model, confirming the appropriateness of the animal model. On the other hand, AAV-SaCas9-KKH-sgRNA significantly increased the proportion of the remaining TH-positive cells in the ipsilateral SNpc to 63.8%, whereas AAV-SaCas9-KKH without sgRNA did not (Figure 4A, C). These findings together indicate that significant A53T-SNCA gene deletion by CRISPR/Cas9 *in vivo* prevents dopaminergic neuronal death in a PD animal model.

### AAV-CRISPR-A53T prevents α-synuclein overexpression-induced forelimb akinesia

We further investigated whether gene editing of A53T-SNCA prevents A53T-SNCA overexpression-induced parkinsonian motor symptoms *in vivo* (Figure 5A). To assess the PD-like motor symptoms, we performed a stepping test, which has been validated in previous studies,^18, 33^ during the second, third, and fourth week after virus injection. As expected, there was no significant difference in the stepping numbers of ipsilateral forelimbs between the groups. On the other hand, stepping behavior of the contralateral forelimb was significantly impaired in the A53T rat model. We found that stepping deficits of the contralateral forelimb was significantly alleviated by AAV-SaCas9-KKH-sgRNA injection, whereas AAV-SaCas9-KKH without sgRNA only had a marginal effect. The ratio of stepping numbers of the A53T KO group was also significantly higher than that of the CRISPR-non-treated and negative control groups (Figure 5D). Taken together, these findings indicate that CRISPR/Cas9-mediated genome editing alleviates A53T-SNCA-induced PD pathology and motor deficits.

**Figure 5.**
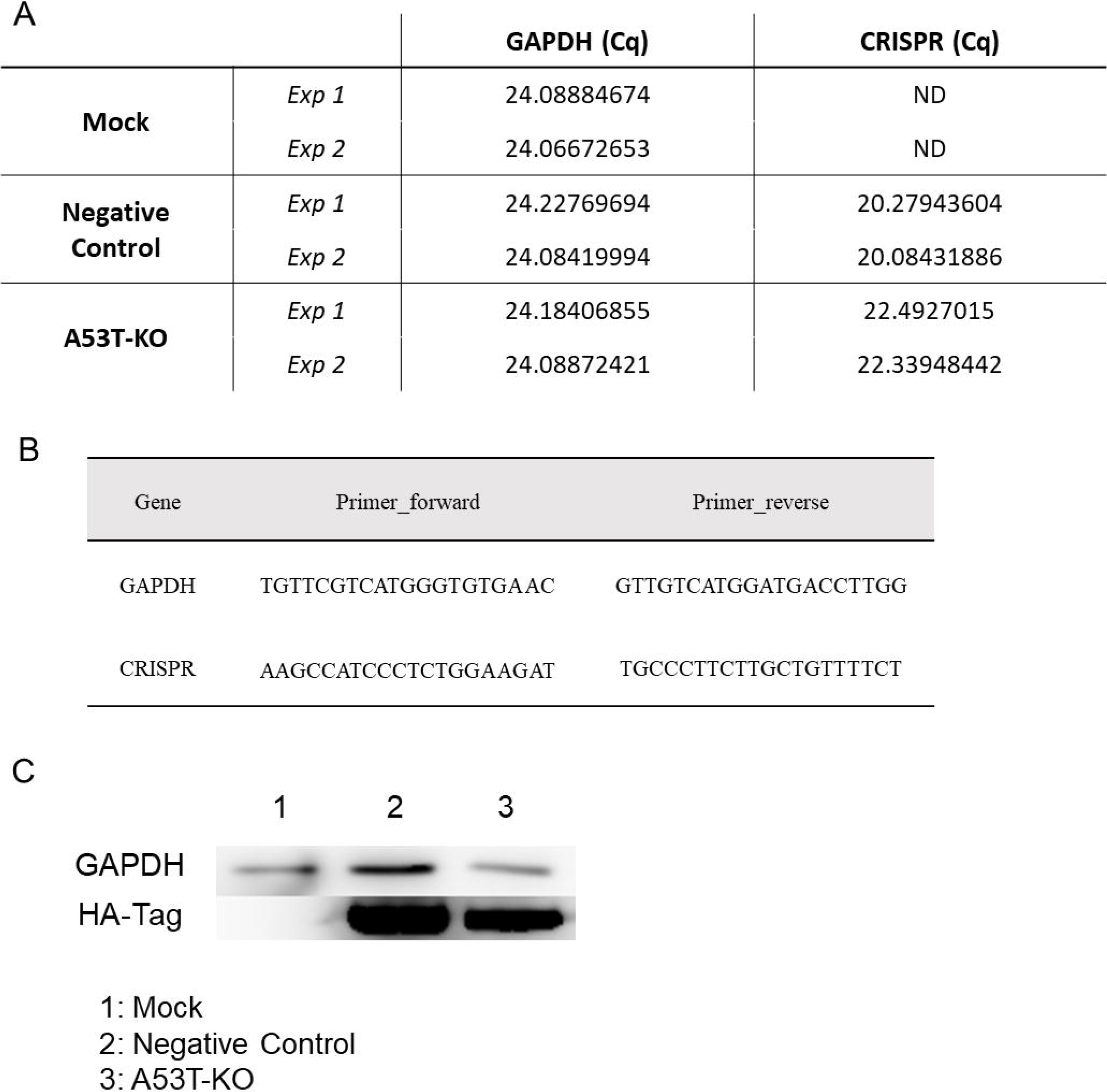
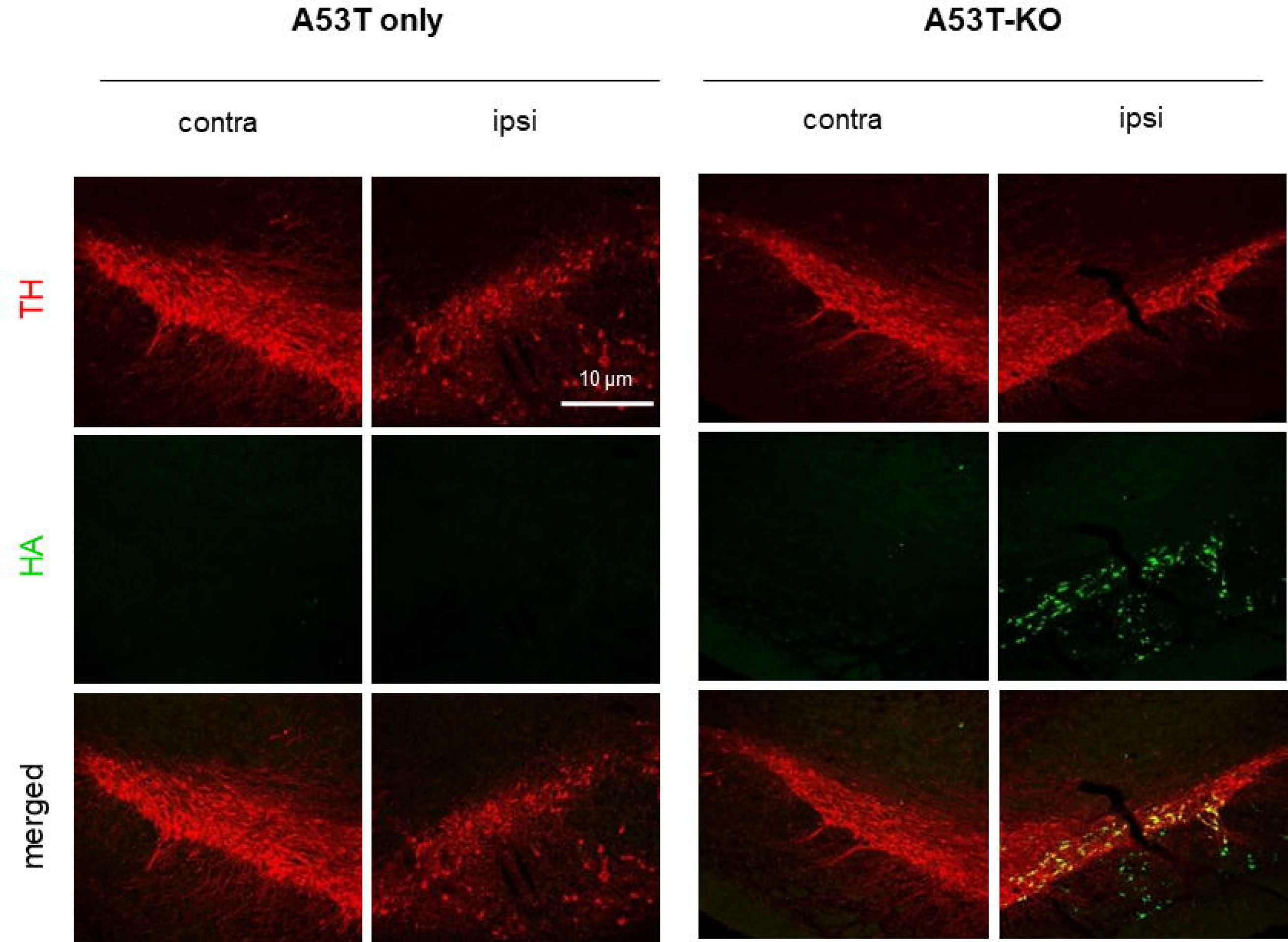
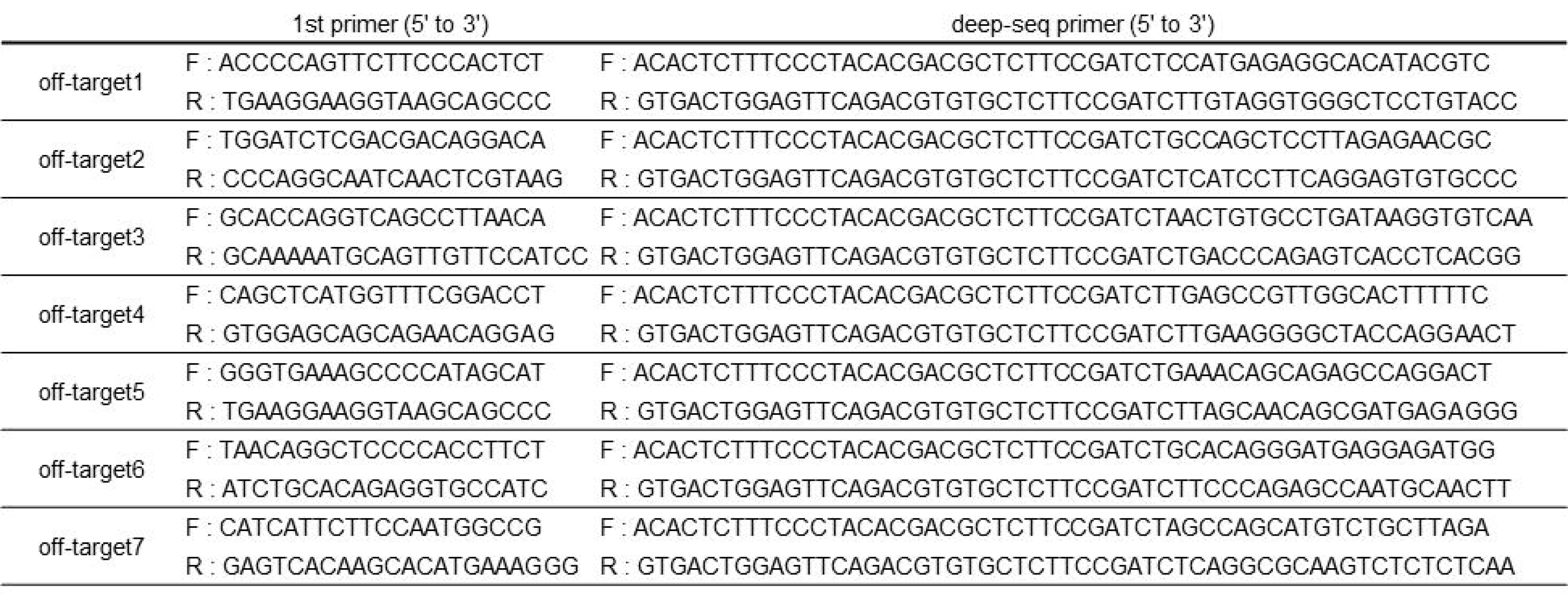
CRISPR-mediated A53T deletion significantly prevents A53T-SNCA-induced parkinsonian motor deficits. (A) Left, schematic diagram of viral injection of AAV-A53T-SNCA simultaneously with AAV-SaCas9-KKH-sgRNA or AAV-SaCas9-KKH without sgRNA into unilateral SNpc. Right, experimental timeline for virus injection and stepping tests. (B) Quantification of adjusted step numbers of the ipsilateral forelimb from the stepping test. (C) Quantification of adjusted step numbers of the contralateral forelimb from the stepping test. (D) Ratio of the adjusted step numbers of the contralateral forelimb over total step numbers. Mean ± SEM. *P < 0.05; **P < 0.01; ***P < 0.001; ns, non-significant (twoway ANOVA with Bonferroni’s multiple comparison test).

### Off-target issues were resolved in the rat brain

The off-target candidates for SaCas9-KKH modification were screened in the whole genome of *Rattus norvegicus* and seven off-target sequences with the highest similarity were chosen (Table 1). Primers for the off-target sequences were designed (Supplementary table 1), and targeted deep sequencing was performed with next generation sequencing (NGS). Off-target sequences were designed from endogenous rat genome, and InDel mutations (%) were analyzed by CRISPR RGEN Tools Cas-Analyzer software (http://www.rgenome.net/). Each group (each n=4, total n=12) showed a low InDel mutation rate (almost less than 1%) and there was no significant difference between the A53T KO group and the other groups. These results indicate that the specificity of SaCas9-KKH-sgRNA targeting A53T-SNCA is very high, which can eliminate off-target issues (Table 2).

**Table 1.**
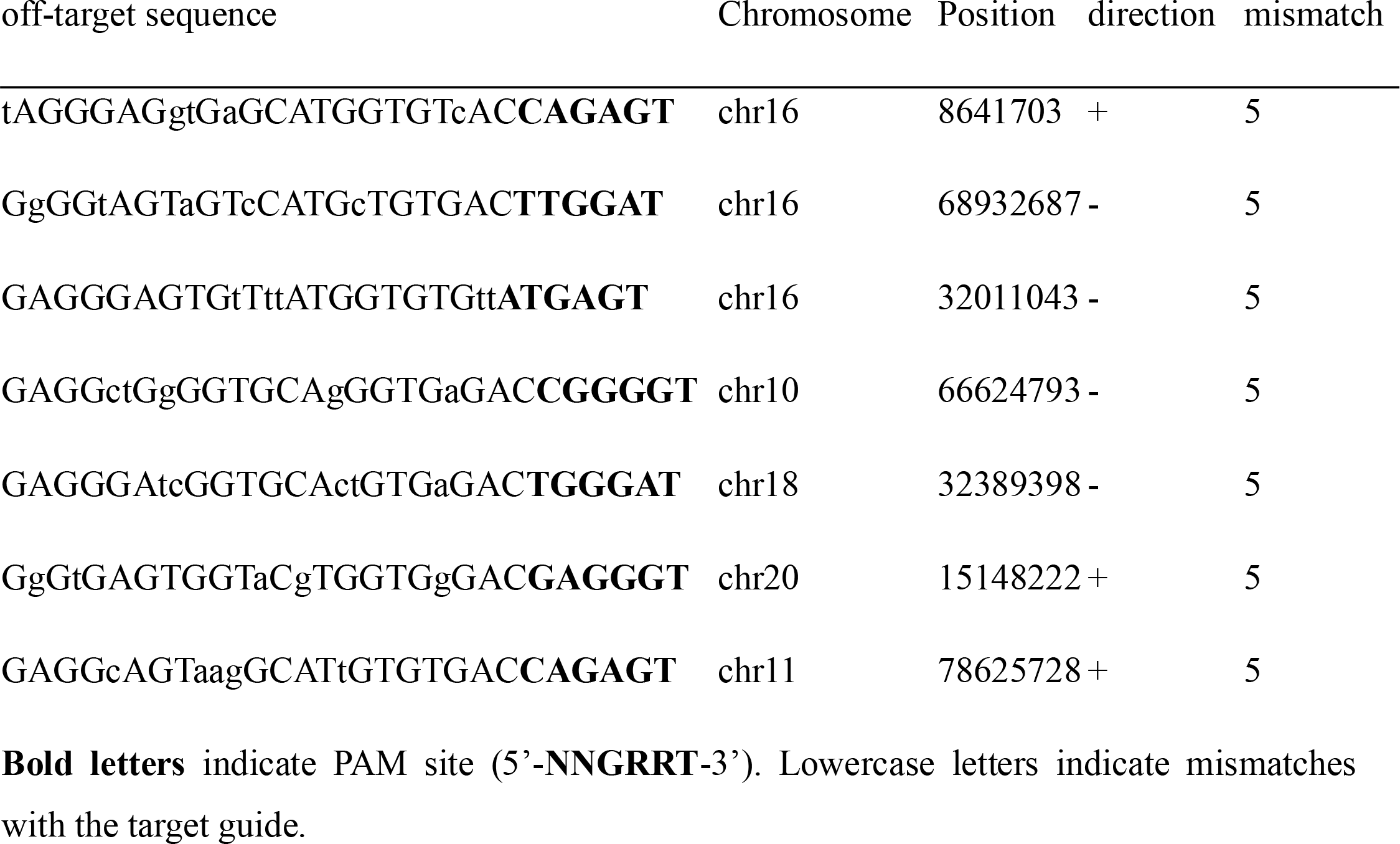
The SNCA off-target sequences of SaCas9-KKH.

**Table 2.**
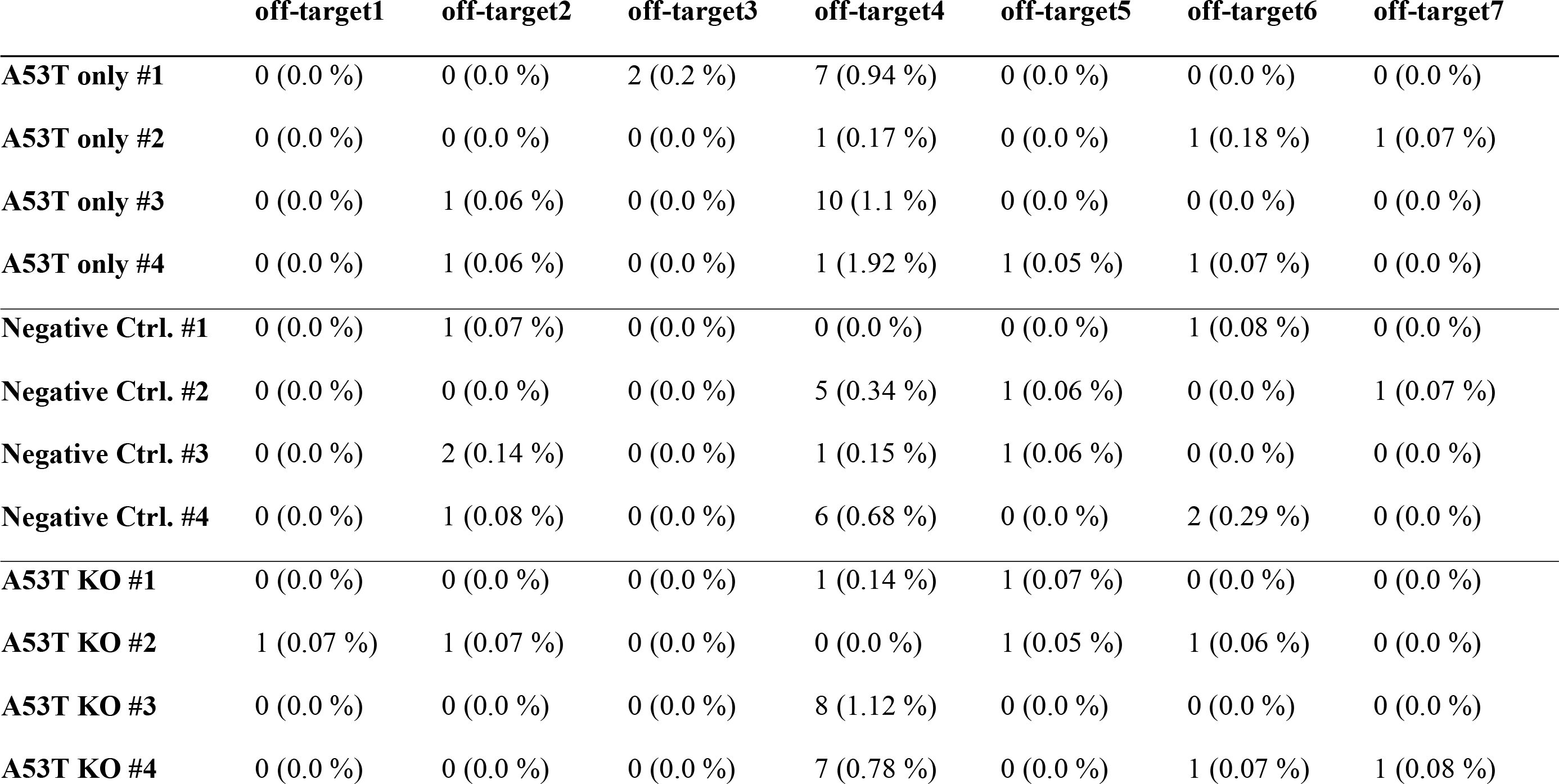
Percentage of off-target InDel mutations.

## Discussion

PD has been thought to be caused by dopaminergic neuronal degeneration in the SNpc, which is strongly associated with the appearance of abnormal cytoplasmic aggregation of mis-folded proteins called Lewy bodies.^1^ A Lewy body is mainly composed of aggregated α-synuclein.^1^ Several mutations of α-synuclein have been reported to accelerate this aberrant aggregation.^10, 34, 35^ Among them, the A53T mutation has been linked to early-onset PD.^8^ The A53T mutation was first documented in families of Italian and Greek descent,^8^ and also documented in Korean^36^ and Swedish^37^ familial cases and a Polish sporadic case.^38^ However, no therapy directly and specifically targeting the A53T-mutated SNCA gene has been developed. In the current study, we designed a gene editing technique for A53T-SNCA using CRISPR and demonstrated that gene deletion of A53T-SNCA can ameliorate forelimb akinesia in an A53T-SNCA-overexpressed mouse model of PD through the cessation of cell deterioration.

In physiological conditions, α-synuclein plays a crucial role in synaptic vesicle trafficking and recycling in presynaptic terminals.^39^ Even though overexpression of α-synuclein leads to substantially worse conditions than those caused by loss of α-synuclein, the genetic deletion of α-synuclein has been reported to cause a reduction in nigrostriatal DA release and an attenuation of DA-dependent locomotor responses to amphetamine.^40^ Moreover, α-synuclein knock-out mice exhibited significant impairments in synaptic response to a prolonged train of repetitive stimulation.^41^ These previous findings suggest that α-synuclein is an essential presynaptic, activity-dependent regulator of synaptic transmission, especially for DA neurotransmission. Therefore, the selective gene deletion of A53T-mutated SNCA, which does not interfere the expression of WT-SNCA gene and the physiological function of α-synuclein, is necessary for alleviating the PD pathology associated with the mutation. In this regard, our finding that CRISPR-mediated gene deletion of A53T-mutated SNCA does not disrupt the expression of WT-SNCA gene implicates CRISPR therapy as a possible strategy for clinical applications.

Since 2013, when CRISPR’s mammalian cell editing function was revealed, interest in overcoming genetic diseases has exploded.^19, 20^ However, even several years later, very few cases of successful application in real diseases have been reported.^32, 42^ The reason for this lack of success is not only the typical clinical trial entry hurdle. Even in animal models such as mice and rats, there are very few examples that simultaneously provide both the experimental evidence that the appropriate level of editing was performed in the tissue and that the behavior was improved. As such, there are important reasons why CRISPR is not easy to be applied in clinical practice.

First, while it is easy to break a target nucleotide sequence using CRISPR, it is not easy to convert it into a specific, desirable nucleotide sequence. The CRISPR obtained in nature cuts the target gene and generates KOs very efficiently through a non-homologous end joining (NHEJ) pathway mechanism. Therefore, it is no exaggeration to say that it has opened a new era in KO cell line and animal model construction technology through cell or zygote KOs. However, the genetic diseases encountered in actual clinical practice very often require an accurate restoration of a broken gene, rather than being solved by simply destroying the function of the gene. To this end, bioengineering using homology-directed repair (HDR) is continuously being developed, but its efficiency and accuracy are still low, so it is difficult to apply it in clinical practice.^32^ To overcome this, Liu opened the way to selectively change only one base by creating groundbreaking CRISPRs called base editors.^43, 44^ However, it is still not ready to be applied in clinical practice because of its low efficiency, accuracy, and expandability.^32^ Therefore, finding a good clinical target that can utilize the superior KO efficiency of CRISPR is crucial. In this study, the original function of CRISPR was adequately utilized by targeting a disease that can be corrected by selective KO of the hotspot mutant gene, A53T-SNCA.

Second, there are several challenges with CRISPR delivery. The method of delivering CRISPR to cells has seen recent advances, and issues with delivery have been resolved to some extent. In particular, the method of delivery of RNP by electroporation is efficient and useful. On the other hand, delivering the CRISPR system to living organisms, even to specific locations or tissue, is still a significant challenge for scientists. Among the various methods, AAV is generally accepted as a relatively safe and one of the most efficient delivery tools in PD patient treatment.^45^ However, even in this case, because of the limitation of cargo size, it is difficult to fully mount SpCas9, which is efficient; therefore, SaCas9 is currently used as the best alternative. SaCas9 is significantly less efficient than SpCas9, and KO efficiency in tissue is usually very low.^32^ For this reason, various KO models using CRISPR have been proposed, but in practice, KO efficiency is low and histological/behavioral results are poor.^42^ However, our study obtained positive data in KO efficiency in histological and behavioral results by meticulous optimization. In particular, CRISPR expression and KO efficiency were sequentially confirmed in all processes; *in vitro*, in-cell, and *in vivo*.

Third, there are potential off-target issues. The CRISPR/Cas9 system detects the desired target accurately and performs double strand breakage. However, CRISPR/Cas9 occasionally also breaks unwanted nucleotide sequences, although this occurs in only a low percentage of cases. This is referred to as an off-target event, and in all animal models, these off-target events must be validated to ensure safety.^46^ The best way to avoid this issue is to design gRNA to target regions where the corresponding nucleotide sequence in the species overlaps as minimally as possible with the nucleotide sequence of another gene. This is referred to as a mismatch, and in general, if there are more than three mismatches, the off-target event is significantly lowered. If inevitably there are only one or two mismatches, designing that location to the seed region (close to the PAM site) is an efficient way to reduce off-target events.^47^ In this study, we selected a location where overlap with other genes was minimal, confirming through BLAST that there were no 0~4 mismatch sequences in *Rattus norvegicus*. In addition, we succeeded in validating rat SNpc tissue using NGS, ensuring that no off-target event occurred in five mismatch genes. Inevitably, there was one mismatch between A53T-SNCA and WT-SNCA (ATA for A53T vs. GCA for WT). We designed sgRNA to be arranged in the second position from PAM (a strong seed region) so that off-target events would occur as rarely as possible. Additionally, we proved its safety through various experiments.

CRISPR/Cas9 is obviously a powerful biotechnological tool, but there are few actual cases showing therapeutic effects *in vivo*. ^28, 29, 32, 48^ In particular, in the field of PD, almost all previous studies presented the possibility of treatment only at the cellular level.^49, 50^ A few papers suggested PD animal model treatment methods; however, all of them applied CRISPR to individual cells as an *ex vivo* treatment technique.^51, 52^ Invariably, there are no reports of successful *in vivo* treatments by delivering CRISPR directly to animals with PD. Our study overcomes several difficulties and demonstrates the possibility of familial PD treatment for the first time in an animal model applying CRISPR to knock out the A53T-SNCA gene *in vivo*, the direct cause of familial PD. This is of great significance in that it has explored the possibility of clinical trials for the development of therapeutic agents using CRISPR to treat PD.

We have assessed the therapeutic effect of CRISPR specifically targeting A53T-SNCA in a PD mouse model. However, the majority of the PD patients suffer from idiopathic PD in which aberrant accumulation of Lewy bodies appear without the A53T mutation of the SNCA gene. Therefore, expression of the WT-SNCA gene should be downregulated, and the WT-SNCA gene should not be knocked out. Because CRISPR itself can only delete the WT-SNCA gene in CRISPR-expressing cells, whether and how the *in vivo* transduction level of AAV conveying CRISPR can be controlled might be an interesting topic that awaits future investigations.

In conclusion, *in vivo* gene deletion of A53T-SNCA using CRISPR/Cas9 significantly protected against α-synuclein accumulation, dopaminergic neurodegeneration, and parkinsonian motor deficits in an A53T-SNCA-overexpressing mouse model of PD. Based on our findings, we propose CRISPR/Cas9 as a potential preventive and/or therapeutic strategy for familial PD with A53T mutation.

## Methods

### pET28a-SaCas9-KKH plasmid construction and protein purification

The SaCas9-KKH gene was amplified by PCR using Q5 polymerase (New England Biolabs, Ipswich, MA, USA) and inserted into the pET28a plasmid between the EcoRI and XhoI restriction sites using 2X HiFi DNA Assembly Master Mix (New England Biolabs) to create pET28a-SaCas9-KKH. After transformation, bacterial cells were grown in 400 mL of Luria broth supplemented with 50 μg/mL kanamycin at 37°C, and 1 mM isopropyl β-D-thiogalactoside (IPTG) was added when the optical density at 600 nm (OD_600_) was 0.4. The cells were grown at 18°C overnight and harvested by centrifugation at 4,000Xg for 10 min, followed by resuspension and sonication in lysis buffer (50 mM Na_2_PO_4_ (pH 8.0), 300 mM NaCl, 10 mM imidazole) supplemented with 1 mM dithiothreitol (DTT), 1 mM phenylmethylsulfonyl fluoride (PMSF), and 1 mM lysozyme. After centrifugation at 4,000Xg for 30 min, the supernatant was recovered and reacted with Ni^2+^-NTA beads for 30 min. Thereafter, the supernatant was removed and the beads were washed three times with wash buffer (50 mM Na_2_PO_4_ (pH 8.0), 300 mM NaCl, 20 mM imidazole), followed by incubation with 5 mL elution buffer (50 mM Na_2_PO_4_ (pH 8.0), 300 mM NaCl, 250 mM imidazole) for 10 min in ice. The SaCas9-KKH protein eluted from the beads was replaced into a conical tube, and then concentrated with a 100K amicon filter (Sigma Aldrich, St. Louis, MO, USA) by centrifugation at 14,000Xg and stored in storage buffer (50 mM HEPES (pH 7.5), 200 mM NaCl, 40% Glycerol 40%, and 1 mM DTT).

### CRISPR sgRNA synthesis

Computationally designed sgRNAs were synthesized by *in vitro* transcription. The sgRNAs were transcribed by T7 RNA polymerase in 40 mM Tris-HCl [pH 7.9], 6 mM MgCl_2_, 10 mM DTT, 10 mM NaCl, 2 mM spermidine, NTPs, and an RNase inhibitor. The reaction mixture was incubated at 37°C for 8 h. The sgRNAs were purified using PCR purification kits (GeneAll, Seoul, Korea) and quantified using a NanoDrop spectrophotometer.

### *In vitro* cleavage assay

The oligonucleotide templates for the *in vitro* SaCas9-KKH cleavage assay, including α-Synuclein wild-type and A53T mutant cDNA, was amplified to 3,000 bp lengths with Sun PCR blend polymerase (Sun Genetics, Daejeon, Korea). The guide RNA was synthesized by *in vitro* transcription with T7 RNA polymerase (New England Biolabs). The oligonucleotide templates (10 ng/μL), SaCas9 (50 ng/μL), and gRNA (25 ng/μL) were mixed in cleavage buffer containing 1.6 mM potassium acetate, 0.625 mM Tris-acetate, 0.31 mM magnesium acetate, and 3.1 μg/mL BSA in pH 7.9. The samples were incubated at 37°C for 20 min and analyzed by agarose gel electrophoresis.

### Plasmid construction

A single vector AAV-Cas9 system containing SaCas9 and its sgRNA scaffold with a CMV promotor was purchased from Addgene (Catalog #61591).^27^ Three sequences of SaCas9 were replaced with E782K, N968K, and R1015H using QuikChange II Site-Directed Mutagenesis Kit (Stratagene, La Jolla, CA) following manufacturer’s instructions to construct pAAV-SaCas9-KKH without sgRNA as a negative control. Subsequently, we additionally designed and cloned a sequence of sgRNA, targeting A53T-SNCA (pAAV-SaCas9-KKH-sgRNA) but not WT-SNCA.

### CRISPR transfection in-cell and cleavage assay using FACS

We tested in-cell whether pAAV-CRISPR-A53T results in sufficient DNA breakage in pAAV-A53T-SNCA-GFP without off-target effects in pAAV-WT-SNCA-GFP. While pAAV-A53T-SNCA-GFP and pAAV-WT-SNCA-GFP are plasmids, we analyzed with flow cytometry whether GFP had been turned off or not. The HEK 293T cell line (HEK) was cultured with Dulbecco’s modified Eagle’s medium (DMEM; Welgene, Gyeongsansi, Korea) containing 10% fetal bovine serum (FBS; Welgene). Identical assays were performed for pAAV-A53T-SNCA-GFP and pAAV-WT-SNCA-GFP, respectively. Three groups were prepared in 24 □well plate with 1×10^5^ cells/well. The first group (mock) was not transfected with any vectors. The second group was used as a negative control, transfected with pAAV-A53T-SNCA-GFP (or pAAV-WT-SNCA-GFP) (50 ng/well) and pAAV-SaCas9-KKH (1 μg/well). The last group was A53T KO, co-transfected with both pAAV-A53T-SNCA-GFP (or pAAV-WT-SNCA-GFP) (50 ng/well) and pAAV-SaCas9-KKH-sgRNA (1 μg/well). Transfection was conducted with Lipofectamine 2000 (Life Technologies, Carlsbad, CA) and Opti-MEM (Life Technologies, Gaithersburg, MD) according to the manufacturer’s instructions and incubated for 72 h. The cells were washed with phosphate □ buffered saline (PBS; Welgene) and detached with Trypsin-EDTA Solution (Welgene), analyzed immediately with Fluorescence-activated cell sorting (FACS, BD FACSCanto II, BD Biosciences, San Jose, CA) system (Figure 3).

### CRISPR gene expression analysis using qRT-PCR

Total RNA was extracted from the cells with RNeasy mini kit (Qiagen) according to the manufacturer’s instructions. Extracted RNA was reverse transcribed to cDNA with Transcriptor First Strand cDNA Synthesis Kit (Roche, Basel, Switzerland). Real-time PCR was performed with a LightCycler^®^ 480 II and a LightCycler^®^480 SYBR Green I Master (Roche, Basel, Switzerland). The relative expression level of the CRISPR mRNA was standardized using GAPDH expression. Results and mRNA primer sequences are provided in the Supplementary Data (Figure S1).

### Western blot analysis

Cells were lysed using RIPA buffer (sigma) with a phosphatase inhibitor cocktail (GenDEPOT). The concentration of extracted proteins was determined using NanoDrop One (Thermo Scientific, Waltham, MA, USA). Samples were heated at 95°C for 5mins and cooled down with ice. Each 80 μg protein was electrophoresed on 4–12% SDS-polyacrylamide gel (Invitrogen, Basel, Switzerland). The gel was transferred onto PVDF membrane. Blots were blocked with PBS containing 0.05% Tween 20 and 5% nonfat dry milk for 2 h, then incubated with GAPDH (Santa Cruz, CA, USA) or HA-Tag (Invitrogen) antibody for 1 h at room temperature. After washing with PBS containing 0.05% Tween 20, blots were incubated with peroxidase-conjugated secondary antibody for 1 h at room temperature. Finally, blots were visualized by enhanced chemiluminescence procedures (Bio-rad, Hercules, CA) according to the manufacturer’s instructions (Figure S1).

### AAV construction

Three previously generated virus vectors were pseudotyped, where the transgene of interest was flanked by inverted terminal repeats of the AAV2 packaged in an AAV-DJ capsid. AAV-DJ was engineered via DNA family shuffling technology, which created a hybrid capsid from AAV serotype 8. AAV-A53T-SNCA, AAV-SaCas9-KKH and AAV-SaCas9-KKH-sgRNA vectors were thereafter purified by iodixanol gradient centrifugation, and Amicon filtration by the KIST Virus Facility. Genomic titers were 2.4 × 10^13^ genome copies /mL (GC/mL) for AAV-A53T-SNCA, 1.5 × 10^13^ GC/mL for AAV-SaCas9-KKH, and 1.8 × 10^13^ GC/mL for AAV-SaCas9-KKH-sgRNA.

### Experimental animals

Twenty-three male Wistar rats (Orient Bio Inc., Seongnam, Korea), weighing 300–350 g at the beginning of the experiment, were housed in a room with a 12-h light/dark cycle and had free access to food and water. All procedures complied with the guidelines of the Institutional Animal Care and Use Committee (IACUC) of the Asan Institute for Life Sciences and were approved by the Ethics Committee for Animal Experiments of the Asan Institute for Life Sciences (Seoul, Korea).

### Stereotactic AAV injection and experimental group

KIST (Korea Institute of Science and Technology, Seoul) virus facility manufactured all AAV vector plasmids. The solution contained a minimum number of viral particles of 1.0 × 10^10^/mL, which is considered a concentrated virus package. Surgical procedures were performed under general anesthesia induced by an intraperitoneal injection of a mixture of 35 mg/kg of Zoletil and 5 mg/kg of Rompun. The AAV vector plasmid was unilaterally injected into the right substantia nigra at the following coordinates: AP −5.4 mm, L +2.0 mm relative to bregma, and V −7.5 mm from the dura. The AAVs were delivered at a rate of 0.2 μL/min using a 33-gauge Hamilton syringe and an automated microsyringe pump (Harvard Apparatus, Holliston, MA, USA). After injection, the needle was kept in place for 5 min to prevent the solution from flowing backward and was retracted over the subsequent 5 min. Twenty-three rats were separated into three groups: 1) the A53T only group (n=7) as a control group, in which each rat was subjected to a virus injection of a mixture of 2 μL of AAV-A53T-SNCA and 1 μL of PBS; 2) the negative control group (n=8), in which each rat was subjected to a virus injection of a mixture of 2 μL of AAV-A53T-SNCA and 1 μL of AAV-SaCas9-KKH; and 3) the A53T KO group (n=8) as an experimental group, in which each rat was injected with a mixture of 2 μL of AAV-SNCA A53T and 1 μL of AAV-SaCas9-KKH-sgRNA.

### Stepping test

All rats were subjected to stepping tests at 2, 3, and 4 weeks after AAV injections. The stepping test was performed as previously described, with slight modifications. Briefly, both hindlimbs were firmly held in one hand of the experimenter, whereas one of the forelimbs was held in the other hand. The test was repeated with both the contralateral and ipsilateral forelimbs. The rostral part of the rat was lowered onto a treadmill (Jeung Do Bio & Plant Co., Seoul, Korea) that moved at a rate of 1.8 m/10 s. The rat’s body remained stationary while one forelimb was allowed to spontaneously touch the moving treadmill track for 10 s. All experimental sessions were video-recorded to allow counting of the number of adjusted steps. Every rat was subjected to the stepping test twice in each session, and the number of steps taken was averaged across the two trials.

### Immunohistochemistry

For tissue fixation, rats were transcardially perfused with 0.9% saline containing 10,000 IU heparin (Hanlim Pharm, Seoul, Korea), followed by 4% paraformaldehyde in phosphate-buffered saline (PBS). Brains were extracted and post-fixed for 12 h in the same fixative, followed by dehydration in 30% sucrose until they sank. Coronal sections (30 μm thick) of substantia nigra (AP, −4.8 to −6.0 mm) were collected using a cryotome (Thermo Scientific, Waltham, MA, USA). The sections were blocked in 0.1 M PBS containing 0.3% Triton X-100 (Sigma), 2% goat serum (ab7481, Abcam) and 2% donkey serum (GTX27475, Genetex) for 1.5 h at room temperature. Primary antibodies used are as follows: chicken anti-GFAP (1:500, ab5541, Millipore), mouse anti α-synuclein (1:300, ab1903, Abcam), rabbit anti-TH (1:500, P40101-0, Pel-freez). Brain sections were incubated in a mixture of primary antibodies overnight at 4 □. After three 5-min washes in 0.1 M PBS at room temperature, the brain slices were incubated in appropriate secondary antibodies from the Jackson Laboratory for 1.5 h at room temperature. Finally, the sections were again washed three times in 0.1 M PBS, then had coverslips mounted on top using fluorescent mounting medium (S3023, Dako). A series of fluorescent images were obtained with an A1 Nikon confocal microscope, and Z-stack images in 2-μm steps were processed for further analysis using the NIS-Elements (Nikon, Japan) software and ImageJ (NIH, MD, USA). Any alterations in brightness or contrast were equally applied to the entire image set.

### Off-target screening of SaCas9

Rat brain tissue containing substantia nigra pars compacta (SNpc) were fixed with 4% paraformaldehyde and precipitated in 30% sucrose in PBS. Whole brain tissue from 3-grouped rats containing SNpc were coronal sectioned and the genomic DNAs were extracted using a DNeasy^®^ Blood & Tissue Kit (QIAGEN, Valencia, CA, USA) according to the manufacturer’s instructions. The off-target candidates of SaCas9 were screened in whole genome of *Rattus norvegicus* by using the Cas-OFFinder program (http://www.rgenome.net/), and seven off-target sequences with the highest similarity were chosen (Table 1). The primers for the off-target sequences were designed by Primer-BLAST (Supplementary table 1). Serial PCRs were performed for targeted deep sequencing and then a MiniSeq sequencing platform (Illumina, San Diego, CA, USA) was used for next generation sequencing (NGS). The generated FASTQ files were analyzed for detecting insertion and deletion mutations, with CRISPR RGEN Tools Cas-Analyzer software (http://www.rgenome.net/), an online tool for assessing genome editing results using NGS data.

### Image quantification

Quantitative analysis of confocal microscopic images was performed using the ImageJ program (NIH). Region of interests (ROIs) were restricted to SNpc using TH images. To measure α-synuclein intensity in the ROI, we converted α-synuclein images into 8-bit and then measured the intensity. The number of TH-positive cells in ipsilateral and contralateral SNpc were counted manually using the multi-point tool in ImageJ by a blinded researcher. Then, the proportions of ipsilateral TH-positive cells over contralateral TH-positive cells were calculated. Four mice per group were sacrificed, and two to three slices per mouse were stained and analyzed for quantification. Quantification was performed by blinded researchers.

### Statistical analysis

Statistical analyses were performed using Prism 7 (GraphPad Software, Inc.). For comparison of multiple groups, one-way analysis of variance (ANOVA) with Tukey’s multiple comparison test was used. For assessment of change of a group by a certain intervention in multiple groups, the significance of data was assessed by two-way ANOVA with Bonferroni’s multiple comparison test. Data from multiple independent experiments was assumed to have normal variance. Data was assumed to be normally distributed. P < 0.05 was considered to indicate statistical significance throughout the study. The significance level is represented as asterisks (*P < 0.05, **P < 0.01, ***P < 0.001; ns, not significant). All data are presented as mean ± SEM. No statistical method was used to predetermine sample size. Sample sizes were determined empirically based on our previous experiences or upon review of similar experiments in the literature. The numbers of animals used are described in the corresponding figure legends or on each graph. Experimental groups were balanced in terms of animal age, sex, and weight.

## Acknowledgements

The authors declare that they have no conflict of interest. This study was reviewed and approved by the Institutional Animal Care and Use Committee (IACUC) Asan Institute for Life Sciences, Asan Medical Center. The committee abides by the Institute of Laboratory Animal Resources (ILAR) guide (approval no. 2018-02-112). This study was supported by grants from the Basic Science Research Program through the National Research Foundation (NRF) funded by the Korean Ministry of Education, Science and Technology (KR) [NRF-2017R1D1A1B03035760, NRF-2019R1C1C1010602], Korea University, Republic of Korea [K1722461, K1809751, K2014091], and KU Medicine-KIST, Republic of Korea [O1902761] to JWH; KIST institutional program (Project No. 2E30180) through the Korea Institute of Science and Technology funded by the Ministry of Science and ICT of Korea to MHN; and Asan Institute for Life Sciences Grant (2020IL0039) funded by the Asan Medical Center, Seoul, Republic of Korea to SRJ.

## Author contributions

M.-H.N., J.W.H., and S.R.J. initiated and designed the project. H.H.Y, S.Y., S.L., S.-J.O., A.J., H.L., N.-R.K., K.K., B.-J.K., and J.L. performed the experiments. H.H.Y, M.-H.N., and J.W.H. analyzed the data and wrote the manuscript.

